# Cytoskeleton remodeling caused by keratin dysregulation triggers tumor aggressiveness via promoting genomic instability and cellular adaption

**DOI:** 10.1101/2025.11.03.686269

**Authors:** Hsiang-Hao Chuang, Brian Yu-Ting Kuo, Kalpana Sriramadasu, Ming-Tsung Lai, Leo Yen-Ting Lin, Yi-Che Chen, Preetham Kumar Boominathan, Tritium Hwang, Chih-Mei Chen, Cherry Yin-Yi Chang, Jim Jinn-Chyuan Sheu

## Abstract

Cellular architecture depends on keratin intermediate filament as a fundamental component, which offers essential mechanical support to fight environmental stresses. Our previous research demonstrated that keratin fusion variants increase tumor aggressiveness through enhanced cancer stemness in oral squamous cell carcinoma. The proper functioning of keratins plays an essential role in maintaining cell structure while determining cell fate. The present study demonstrates that keratin fusion variant drives genomic instability through cytokinesis defects, which results in the formation of polyaneuploid cancer cells (PACC). The cells expressing keratin fusion show elevated DNA damage repair gene expression, which serves as a key factor for mitotic slippage during cancer development. The PACCs generated by keratin fusion make cancer cells resistant to cisplatin treatment while simultaneously reducing γ-H2AX induction and increasing survival rates. The Gene Set Enrichment Analysis results showed increased “regulation of actin cytoskeleton” activity in keratin fusion-expressing cells correlated with elevated actin filament networks and increased cell motility in these cells. In summary, the keratin fusion variant enhances cancer aggressiveness through three mechanisms: it creates genomic instability that leads to PACC formation and enables cancer cells to evade cGAS/STING-mediated death signals and modifies cytoskeleton structures, which results in drug resistance and metastasis.

## Introduction

Cytoskeleton is composed of networks of microtubules, actin filaments, and intermediate filaments, which work together to preserve the cell’s internal mechanical structures and shape, control transport of intracellular cargo, cell proliferation, cell motility, mechanotransduction, and fate [1]. Research findings show that cytoskeleton network modifications play a vital role in tumor progression and metastasis [2]. The cellular architecture includes keratin intermediate filaments as one of its main components, which offer essential mechanical support to withstand environmental stresses. Keratin dysfunction causes diseases associated with dermatology and even cancers [3–6]. Furthermore, our previous study showed that keratin fusion mutations are associated with poor prognosis, and a mechanistic investigation verified that upregulation of transforming growth factor beta (TGF-β) and granulocyte colony-stimulating factor (G-CSF) signaling contributed to cancer cell stemness, drug tolerance, and cancer aggressiveness [7]. However, we still know relatively little regarding the functions of keratin fusion variants (KFs) in tumor malignancy.

Genomic instability (GIN) is a critical hallmark of almost all cancers, characterized as increased frequency of changes in nucleic acid sequences within the genome, chromosomal rearrangements, or alterations in chromosome numbers (aneuploidy) [8]. Therefore, there are various types of genomic instability in cancer. Chromosomal instability (CIN) is the major form of GIN, harboring alterations in chromosome structure and number. The chromosome cycle and centrosome cycle are normally coupled, and deregulation of the centrosome cycle would lead to chromosomal instability [9]. GIN has been regarded as a driving force of heterogeneity, plasticity, and evolution in tumors [10–12]. Increased evidence shows that genetic variability can drive survival niches such as angiogenesis, metastasis, and drug resistance in cancer [13–16]. Therefore, targeting GIN seems a potential therapeutic strategy for cancer treatment.

Cancer cells with giant or multiple lobes of nucleus containing altered numbers of chromosomes are regarded as polyaneuploid cancer cells (PACCs). The corresponding mechanism of PACCs formation is characterized through cytokinesis failure, mitotic slippage, endo-replication, cell fusion, or cancer cell cannibalism [17]. Initially, this subpopulation has been typically ignored as dormant or senescent, or dying cells upon stress such as chemo- or irradiation-therapies [18]. However, growing evidence has shown that PACCs play an important role in the origin, immortality, invasion, metastasis, and resistance of tumor cells to radiotherapy and chemotherapy, leading to disease progression and relapse [18–21]. It provides a driving force for tumor evolution and malignancy. In our preliminary results, an elevated ratio of PACCs was observed in keratin fusion mutation-expressing cells.

The type I interferon pathway mediated by the cyclic GMP-AMP synthase (cGAS)-stimulator of interferon genes (STING) was initially discovered as an innate immune defense program to early detect foreign pathogen infection, such as cytosolic DNA, and initiate host defense countermeasures [22–24]. In addition to pathogen infections (bacterial, DNA virus), uptake of dying cells, and mitochondria disruption [25], emerging evidence indicates that genomic instability–induced micronucleus formation activates the cGAS–STING pathway, functioning as a cell-intrinsic immune surveillance mechanism [26–30]. Normally, it suppresses tumor progression through the recruitment of immune cells to promote the clearance of tumor cells harboring micronuclei to maintain genome stability or reduce heterogeneity [31, 32]. However, dysregulation of the cGAS-STING pathway would promote tumorigenesis and aggressiveness [33–35].

In the present study, we found that keratin fusion variant promotes chromosome instability, leading to genome instability, accompanied by elevated chemotherapy resistance and cell motility, leading to tumor aggressiveness. Interestingly, keratin fusion variant not only promotes genomic instability but also affects cGAS-STING signaling to remain survival niches under DNA damage stress. It implies that KFs might be a driving force for tumor aggressiveness via the induction of genomic instability and cytoskeleton reorganization.

## Materials and Methods

### Reagents

Detailed information regarding the materials and reagents used in this study is reported in Supplementary Table S1.

### Cell culture

The CAL27 and FaDu human oral squamous cell carcinoma (OSCC) cell lines were purchased from the American Type Culture Collection (ATCC) and the Bioresource Collection and Research Center (BCRC), Taiwan, respectively. Both cell lines were cultured in Dulbecco’s Modified Eagle Medium (DMEM, Gibco, Grand Island, NY, USA) medium supplemented with 10% fetal bovine serum (FBS) (Gibco, Grand Island, NY, USA) and 1X Antibiotic-Antimycotic (Gibco, Grand Island, NY, USA). GFP-fusion protein-expressing cells were maintained in complete medium containing 800 μg/ml G418. mCherry-fusion protein-expressing cells were maintained in complete medium containing 250 μg/ml hygromycin. All cells were maintained at 37°C and 5% CO_2_ in a humidified incubator.

### Immunofluorescent staining

Cells were seeded on 18-mm round coverslips in a 12-well culture plate. Cells were washed and fixed in 4% paraformaldehyde in PBS for 20 min, and permeabilized with 0.5% Triton in PBS for 5 min. After fixation, the cells were subjected to immunofluorescent staining with corresponding primary antibodies for 2 hours, and subsequently probed with Alexa Fluor 488 or 594 secondary antibodies and 1 μg/mL 4′, 6-diamidino-2-phenylindole (DAPI). For γ-H2AX determination, cells were fixed with 2% paraformaldehyde for 20 min and subjected to immunostaining as mentioned above [36]. For gamma-tubulin staining, cells were fixed with cold methanol on ice for 15 minutes, and stained with anti-GFP and anti-γ-tubulin antibodies for 2 hours at room temperature, followed by probed with Alexa Fluor 488 or 594 secondary antibodies and DAPI. For F-actin determination, cells were washed and fixed with pre-warmed solution. The fixed cells were co-stained with Rhodamine-Phalloidin and DAPI to observe F-actin in the cells. The fluorescent images were captured by an inverted epifluorescence microscope (OLYMPUS, Tokyo, Japan).

### Cell cycle analysis

The cells were harvested and washed with PBS buffer, and then fixed in 70% (v/v) ethanol. The fixed cells were stained with propidium iodide solution and injected into a Sony SH800 Cell Sorter (Sony Biotechnology Inc., California, USA) to analyze the cell cycle profile.

### Western blot analysis

The cells were harvested, and the lysates will be prepared in 1× RIPA buffer containing protease inhibitors and phosphatase inhibitors. Protein concentrations were determined with a BCA Protein Assay kit (G-Biosciences) following the manual guidance. The 30 μg of protein lysates were resolved in a SDS-polyacrylamide gel electrophoresis and then transferred onto a PVDF membrane. The specific protein bands were identified and probed with the indicated primary antibodies, followed by horseradish peroxidase (HRP)-conjugated secondary antibodies and displayed by enhanced chemiluminescence solution (ECL, Millipore).

### Transwell migration/invasion assay

Cell migration and invasion ability were determined by using transwell filter inserts (8-μm pore size) and inserts coated with 10% Matrigel (CORNING) for invasion assay, followed by placing them into a 24-well plate (Merck Millipore, Billerica, MA, USA). Briefly, 200 μl of serum-free medium containing 5 × 10^4^ cells was inoculated onto the upper chamber of the transwell. 800 μl complete medium (supplemented with 10% FBS) was loaded onto the lower chamber and incubated for 24 hours at 37°C. Then, the filter inserts were fixed with 4% formaldehyde and permeabilized by using 0.5% Triton X-100. After fixation, the cells were stained with 0.5% crystal violet solution. The filter membranes were washed twice with PBS, and the cells on the top surface of the filter were removed using cotton swabs. The cell numbers were counted from five random fields under a fluorescence microscope (OLYMPUS IX83) at 100X magnification.

### RNA isolation and sequencing

A total of 1 × 10^6^ CAL27 parental and keratin-expressing cells were seeded in 6-well plates in complete media overnight. The next day, the cells were harvested, and total RNA was extracted by using the FairBiotech kit (NRB0100). The extracted RNA was subjected to the preparation of high-quality RNA-seq libraries with KAPA mRNA HyperPrep Kit and sequenced on a Novseq 6000 (Illumina) by TOOLS Biotech Company (Taiwan). Sequenced reads quantified with Trimmomatic were mapped to the human reference genome GRCh38 using HISAT2. We used DESeq2 to normalize gene expression, and DESeq2 R package was also applied to detect differentially expressed genes (DEGs) based on the Poisson distribution model. We then used clusterProfiler to perform Gene Ontology (GO) and KEGG pathway enrichment analysis of DEGs. Additionally, Gene Set Enrichment Analysis (GSEA) was used to identify enriched biological functions and activated pathways from the molecular signatures database (MSigDB).

### EdU incorporation assay

EdU incorporation assay was performed by following the instructions in the manual (Abbkine, Georgia, USA). Briefly, Keratin fusion variant expressing sublines were incubated with 10 μM EdU for 2 h at 37°C. Cells were fixed with 4% formaldehyde for 15 min, and permeabilized with 0.5% Triton X-100 for 15 min at room temperature, followed by incubation with Click-iT reaction mixture for 30 min. DAPI was used to stain with nuclei. The proportion of EdU-labelled cells was determined by a fluorescence microscopy (OLYMPUS IX83).

### Sphere formation

The cells were harvested and diluted to a concentration of 2 × 10^3^/ml in complete medium. 200 μl of cell mixture (4 × 10^2^ cells) was inoculated onto each well of a 96-well low-attachment plate and incubated for 7 days. We analyzed the sphere formation (size bigger than 50 μm) by using a microscope (OLYMPUS IX83).

### Statistical analysis

GraphPad Prism (GraphPad software, La Jolla, CA) was used to analyze the statistical differences by using the two-tailed Student’s t-test or one-way ANOVA for group comparisons. The data in this study are presented as the mean ± standard deviation of at least three independent experiments. * p < 0.05, ** p < 0.01, and *** p <0.001 indicate significant differences among the experimental groups.

## Results

### Keratin acts as a guardian of genomic stability

The functions of the cytoskeleton network are to maintain cell shape, control transport of intracellular cargo, cell cycle progression, cell motility, and mechanotransduction [1]. The protein scaffold linked to its associated complexes forms the main resilience structure. Dysfunction of cytoskeleton members such as keratin can cause diseases associated with dermatology and cancers. Our previous study showed that KFs are associated with poor prognosis, and a mechanistic investigation verified that upregulation of TGF-β and G-CSF signaling contributed to cancer cell stemness and cancer aggressiveness [7]. Furthermore, we also found that tumor tissues composed of cells expressing K6-K14/V7-GFP (**Figure S1A**) harbored enlarged nuclei and pleomorphism [7]. In line with this phenomenon, high ratios of di-nucleus or multi-nucleus in K6-K14/V7-GFP expressing cells were observed. However, the detailed mechanisms of the contribution of KFs to tumor malignancy are still unknown. In the present study, we aimed to decipher the underlying mechanism and the consequences. First, we gained the keratin-expressing clones by G418 selection. We also determined the expression of keratin by Western blotting (**Figure S1B**). Then, we systematically analyzed the nucleus status of keratin-expressing cell sublines. Compared to GFP, K14-GFP expressing cells, V7-GFP expressing ones display a high proportion of nuclear deformity, multi-nucleus, micronucleus, and lagging chromosomes containing metaphase lagging and anaphase bridges in CAL27 cells (**Figure 1A and 1B**). Not only in CAL27 cells, we also found that V7-GFP overexpression dramatically increased the ratios of nuclear deformity, multi-nucleus, and micronucleus formation in FaDu cells (**Figure S2**). It indicates KFs promote genomic instability. Despite multi-nucleus and micronucleus existing, KFs expressing cells showed normal cell cycle progression in flow cytometry analysis (**Figure 1C**). An increased ratio of G2/M phase was observed in V7-GFP expressing cells. Moreover, a higher ratio of S phase was shown in V7-GFP 8-9 cells (**Figure 1C**). Furthermore, we not only found multi-nuclei existing in KFs expressing cells but also in the specimens of KF mutant patients (p=0.0028) (**Figure 1D**).

**Figure 1.**
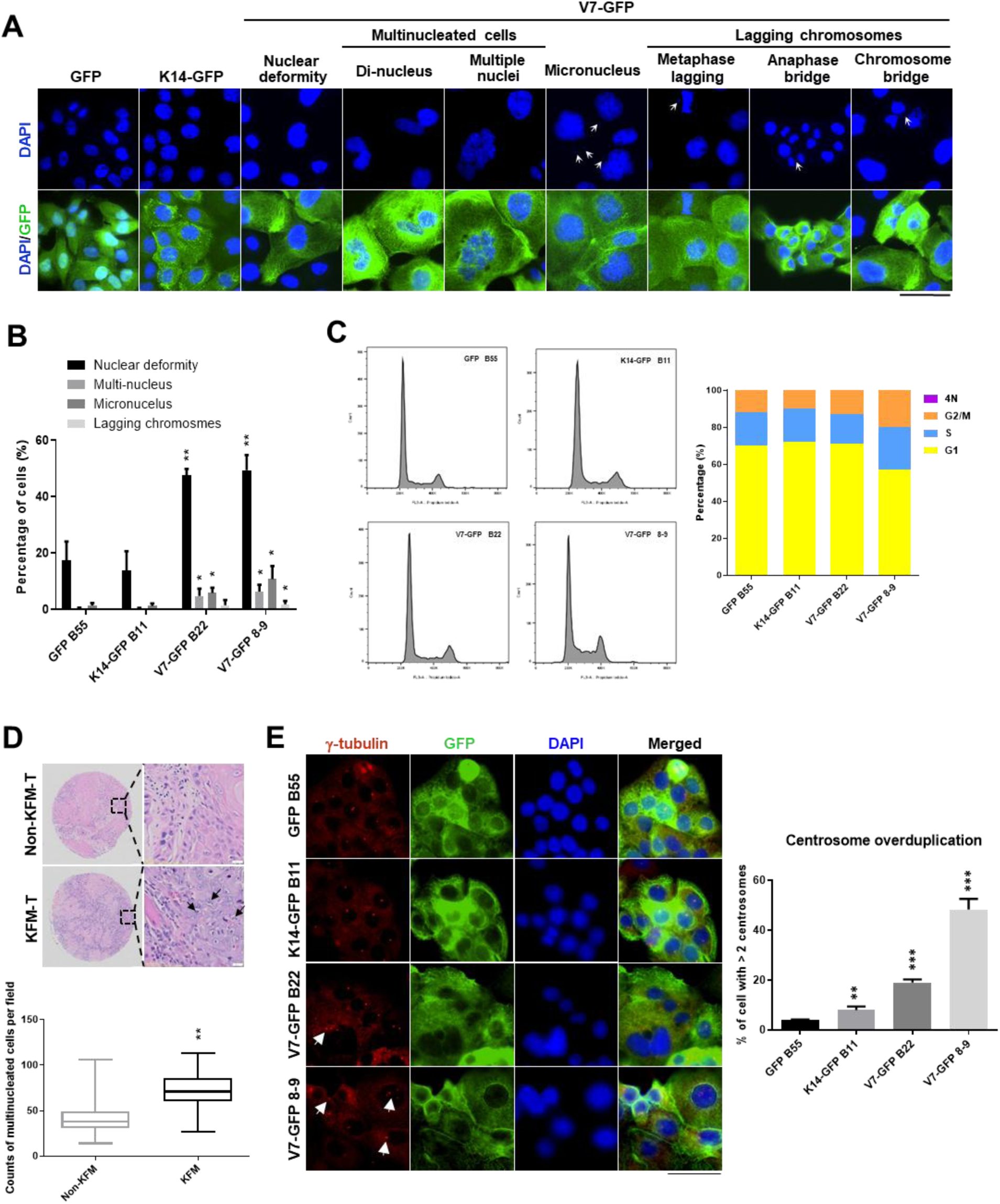
Keratin fusion variant contributes to the genomic instability. (A) CAL27 cells overexpressing GFP, K14-GFP, and V7-GFP were fixed with 4% paraformaldehyde and stained with DAPI. The nuclear status was determined by monitoring DNA staining using fluorescence microscopy. Bar scale indicates 50 μm (B). The nucleus status was counted with the cell_counter (Image J, USA) and plotted in the bar chart. *P < 0.05, **P < 0.01 according to a Student’s t-test. (C) CAL27 cells overexpressing GFP, K14-GFP, and V7-GFP were stained with PI and subjected to cell cycle analysis using flow cytometry (Sony SH800). The percentage of cells in the G0/G1, S, G2/M, S phases of the cell cycle was plotted in the bar chart. (D) The numbers of cells with multi-nuclei in the tissue microarray of oral cancer patients were counted and plotted in the box plot. **P < 0.01. DAPI, 4′,6-diamidino-2 phenylindole; PI, propidium iodide. (E) Keratin-expressing cell sublines were fixed with cold ethanol and stained with GFP, γ-tubulin, and DAPI to determine the centrosome numbers by counting γ-tubulin foci. White arrows indicate centrosome overduplication. The number of centrosomes per cell was scored. More than 100 cells were counted for each experiment group (n=3). Scale bar indicates 50 μm. **P < 0.01 and ***P < 0.001 according to a Student’s t-test.

Multi-nucleus, micronucleus, and lagging chromosomes accompanied by genomic instability are usually caused by defective cytokinesis [37]. Centrosome acts as a microtubule-organizing center (MTOC) [38]. Normally, centrosome duplication is coupled with the chromosome replication cycle to control genome integrity. And centrosome dysregulation usually resulted in cytokinesis defective [39]. Therefore, we determined whether KFM perturbs the function and number of centrosomes by counting γ-tubulin foci in KFM-expressing cells. Most of the GFP and K14-GFP expressing cells harbored a single nucleus per cell. However, V7-GFP expressing cells (B22 and 8-9) exhibited the phenomenon showing more than 2 centrosomes in a cell and accompanied by two or multi-nuclei in a cell (**Figure 1E**). We also observed cytokinesis defective in V7-GFP expressing cells (**Figure 1E**). We further determined the centrosome number to confirm the hypothesis. From the results, we found that most of GFP GFP-expressing control cells have less than 2 centrosomes per cell. K14-GFP expressing cells exhibited a slightly higher ratio of centrosome amplification than control cells. V7-GFP expressing cells harbor a dramatically higher ratio of centrosome overduplication, especially in the 8-9 subline (**Figure 1E**). It indicates that keratin fusion mutation promotes centrosome dysregulation, leading to defective cytokinesis.

### Keratin dysfunction affects the expression and cellular localization of lamins

From our previous study, we found KFs triggered nuclear array dysregulation [7]. And we further determine whether KFs affect the expression or distribution of nuclear lamins, which control nuclear membrane integrity. In the normal condition, most GFP and K14-GFP expressing cells exhibited round or smoothly oval nuclei, as visualized by lamin A/C and DAPI staining (**Figure 2A**). However, distorted or multi-nuclear morphologies were observed in the V7-GFP expressing cells (**Figure 2A**). Interestingly, we found that the expression of lamin A/C in V7-GFP 8-9 CAL27 cells is dispersed (**Figure 2A**). We also observed similar expression patterns of lamin B1 and B2 in keratin-expressing cell lines (**Figure 2B and 2C**). In line with lamin A/C, the expression patterns of lamin B1 and B2 were dispersed in the giant V7-GFP 8-9 cells. It implies that the keratin fusion variant would affect the function of nuclear lamins. In addition to the cellular localization of nuclear lamins, we further determine whether the keratin fusion variant influences the expression levels of nuclear lamins. Compared to GFP and K14-GFP expressing cells, similar levels of lamin A/C were observed in V7-GFP expressing cells, but reduced ones of lamin B1 and lamin B2 were found in V7-GFP expressing ones (**Figure 2D**). It indicates keratin dysregulation affects the structure of the nuclear envelope, leading to alteration of the integrity of the nuclear membrane.

**Figure 2.**
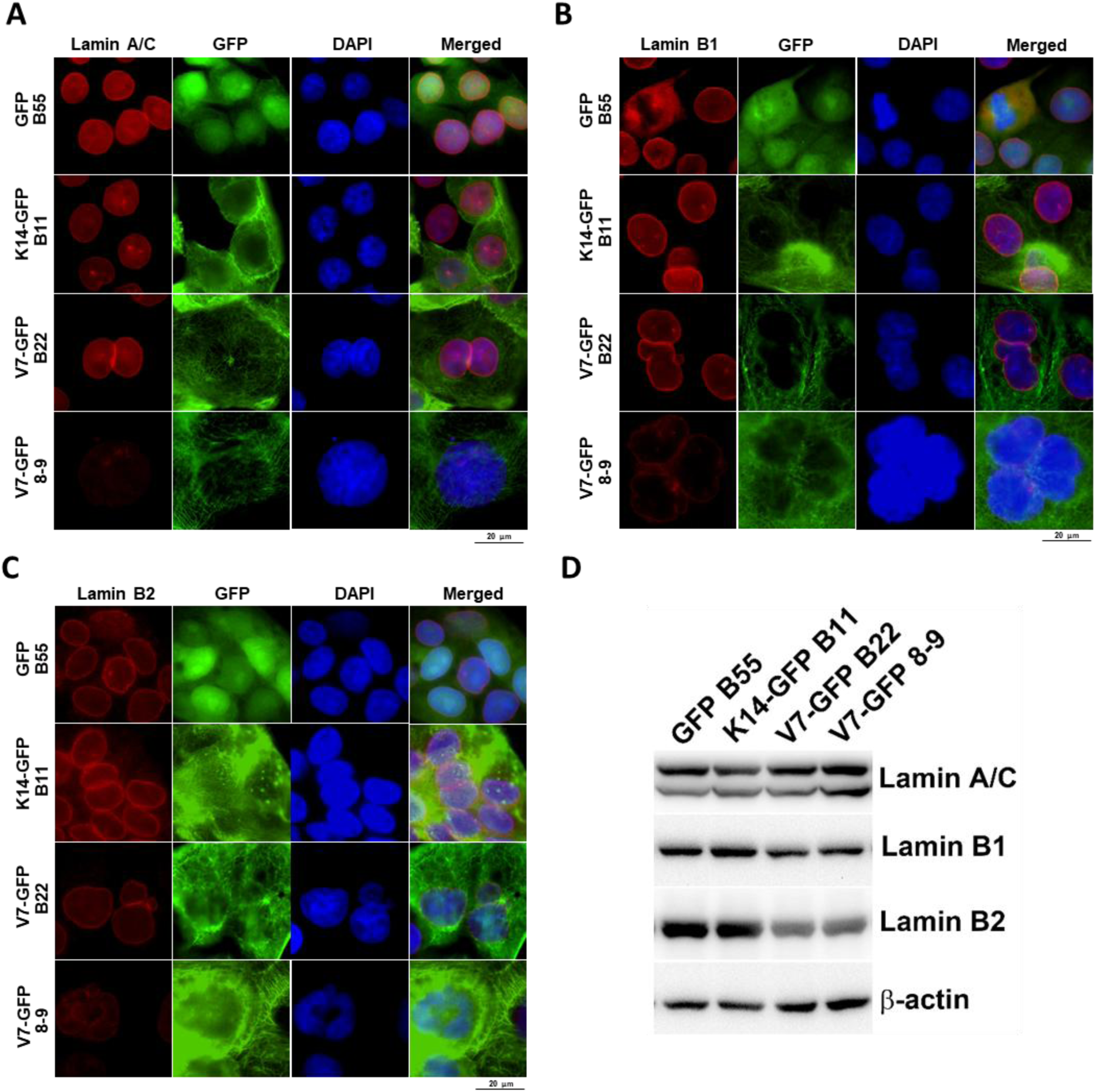
Keratin fusion variant reduces nuclear membrane integrity. Keratin-expressing cell sublines were fixed and stained with lamin A/C (A), lamin B1 (B), and lamin B2 (C), respectively, to visualize the morphology of the nuclear envelope. Nuclei in cells were visualized with DAPI staining. Scale bar indicates 20 μm. (D) The keratin-expressing cell sublines were harvested and lysed and subjected to immunoblotting for the indicated proteins. The level of β-actin was regarded as a loading control.

### Keratin dysfunction manipulates the cGAS-STING signaling axis for survival

In normal conditions, DNA damage response signaling controls genome stability. From previous results mentioned above, keratin fusion variant promotes the genomic instability (**Figure 1**), but cells harboring KFs still proceed normal cell cycle progression (**Figure 1C**). This somehow conflicts with each other. First, we determined whether the keratin fusion variant affected the expression levels of cell cycle-associated molecules. From the results, we found that reduced expression of cyclin-dependent kinases (CDKs) and cyclins but increased expression levels of CDK inhibitors (p16 and p21) in CAL27 expressing V7-GFP B22 cells (**Figure 3A**). This indicates that V7-GFP B22 cells exhibited a dormant-like phenomenon (**Figure 3B**). However, elevated levels of CDKs and cyclins and low expression levels of CDK inhibitors were observed in giant cell-like V7-GFP 8-9 cells (**Figure 3A and 3B**). From previous results, we found that the keratin fusion variant promotes centrosome amplification, leading to genomic instability (**Figure 1**). We further determined whether the keratin fusion variant affected centrosome biogenesis and cytokinesis-associated gene expression. We found that most of these genes were downregulated in V7-GFP B22 cells compared to GFP control cells and K14-GFP expressing ones (**Figure 3A and 3B**). However, these were upregulated in V7-GFP 8-9 cells (**Figure 3A and 3B**). It implies that different clones bearing V7-GFP overexpression trigger genomic instability through distinct pathways. Genomic instability is a critical cancer hallmark resulting from the accumulation of DNA damage [40]. We also determined whether keratin fusion variant influences DNA damage responses-associated genes expression [41, 42]. Compared to GFP and K14-GFP groups, V7-GFP B series cells exhibited reduced levels of DNA damage response-associated genes containing sensors, transducers, mediators, and effectors, except for ones of CDKIs (**Figure 3A and 3B**). Interestingly, giant cell-like V7-GFP expressing cells harbored higher expression of DNA damage response-associated ones, but dramatically reduced levels of CDKIs and STING (**Figure 3A and 3B**). It implies that PACC-like V7-GFP expressing cells harbored higher DNA damage repair machine activity against endogenous damage stress, but dormant-like V7-GFP expressing ones were insensitive to endogenous damage stress. From previous results, KFs trigger micronuclei formation (**Figure 1**), leading to activation of cGAS-STING-mediated interferon responses to eliminate chromosome instability (CIN) [28, 34]. We determined the cGAS and STING expression in keratin fusion-expressing cells harboring micronuclei. The expression of cGAS and Sting 1in V7 B22 cells was observed to increase in micronuclei containing ones but not in V7 8-9 cells (**Figure 3C**). We also characterized the effect of KFs on cGAS-STING-mediated signaling axes. Therefore, we further determined whether KFs affect the cGAS-STING signaling axis. We found that the expression levels of cGAS and STING were increased in V7-GFP B22 cells and accompanied by upregulation of the Type I IFN signaling, such as STAT1 and JNK, meanwhile sustained p38MAPK and STAT3-mediated cell survival signals [35] (**Figure 3D**). However, STAT1 and STAT3 signaling axes were dramatically reduced in giant cell-like V7-GFP 8-9 cells. JNK and p38MAPK-mediated survival signaling was sustained in V7-GFP 8-9 cells [43]. Intriguingly, micronuclei-elicited cGAS-STING signaling were dramatically suppressed in giant cell-like V7-GFP 8-9 cells via downregulating the expression of cGAS and STING (**Figure 3C**).

**Figure 3.**
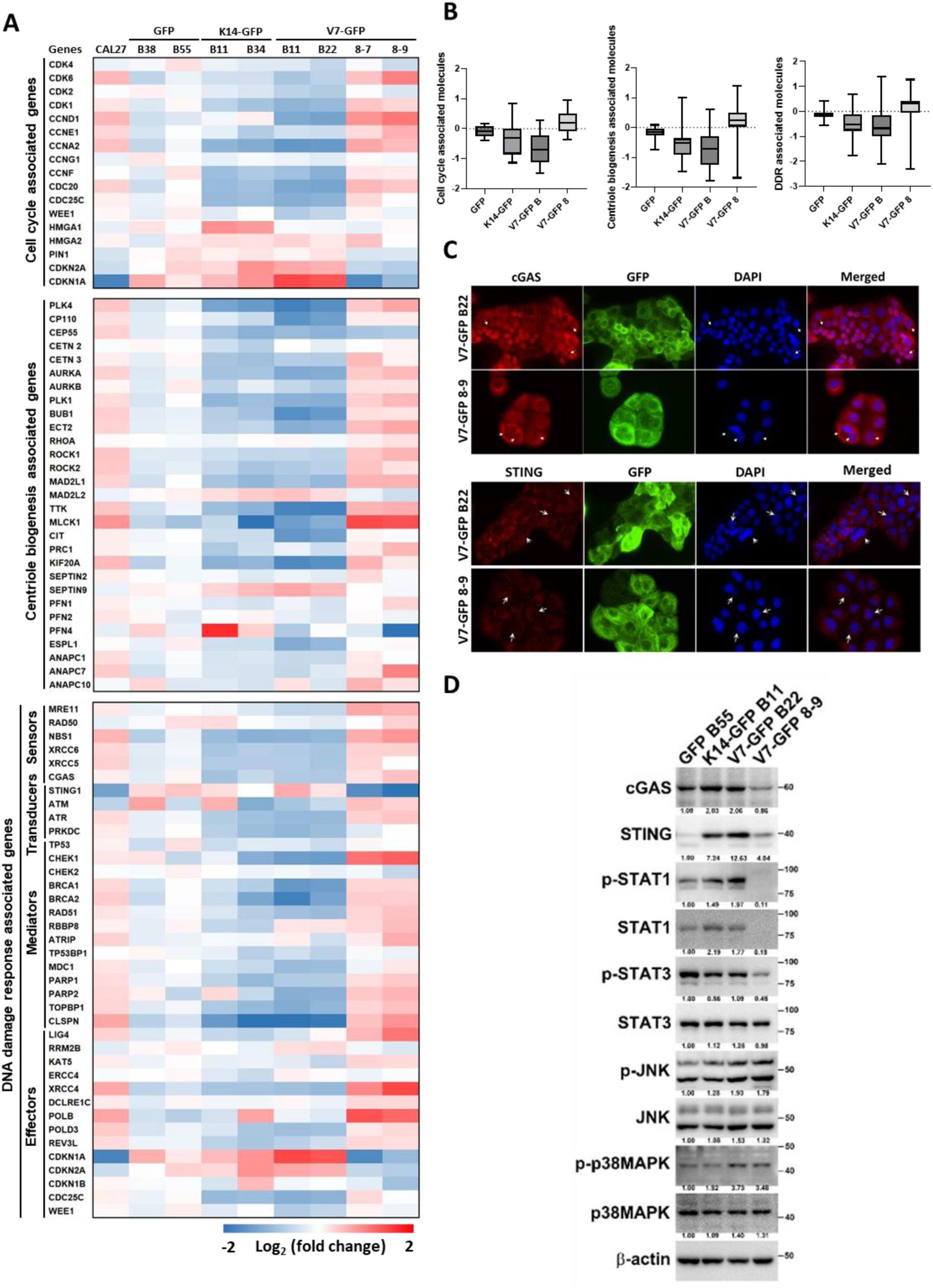
Keratin fusion V7 variant escapes cGAS-STING-mediated cell death signaling via distinctive pathways. Keratin-expressing CAL27 sublines were harvested and extracted the total RNA. The total RNAs were subjected to RNA sequencing. (A) The transcriptional expression levels of cell cycle-associated (upper panel), centrosome biogenesis-associated (middle panel), and DNA damage response-associated molecules (bottom panel) were analyzed in keratin-expressing cell lines. The gene expression is normalized to the mean of parental cells and GFP control ones, and the fold change is calculated with log_2_. (B) The average gene expression with log_2_ calculation in keratin fusion expression cell sublines was drawn in a box plot for cell cycle-associated, centriole biogenesis-associated, and DDR-associated molecules. (C) Keratin fusion expressing cell sublines were fixed and stained with cGAS or STING, and nuclei were visualized with DAPI staining. Arrows indicated micronuclei-containing cells. Scale bar indicates 50 μm. (D) The keratin-expressing cell lines were harvested and lysed. The lysates were subjected to immunoblotting for the indicated proteins. The expression levels of β-actin were used as loading controls. The numbers below the blot images indicate the expression levels relative to the GFP control group and normalized to the β-actin signal intensity.

### Keratin fusion variant increases drug resistance

We found that almost all DNA damage response-associated genes were altered in V7-GFP expressing cells (**Figure 3A**). It implies that keratin fusion variant expressing cells proceed cell cycle progression through repressing DNA damage response signaling under a chromosome instability status. We further determine whether the keratin fusion variant would suppress chemotherapy-induced genotoxic stress (DNA damage stress) to survive under stress conditions. We treated keratin-expressing cell sublines with 10 μM cisplatin (a general chemotherapy drug) for 24 hours and fixed cells to determine the intensity levels of γ-H2AX (DNA damage signaling) using immunofluorescence. We found that either GFP-expressing cells or K14-GFP-expressing ones increased γ-H2AX intensity upon cisplatin treatment (**Figure 4A**). However, V7-GFP expressing cells showed lower levels of γ-H2AX intensity upon cisplatin treatment (**Figure 4A**). It indicates that the keratin fusion variant might suppress chemotherapy-induced genotoxicity-mediated cell death. Due to KFs expressing cells showing dysregulation in DDR expressing profiling, we further determined whether KF variant develops a survival advantage under chemotherapy agent-induced stress conditions compared to control cells. We co-cultured keratin-expressing cells with mCherry-expressing control cells in the absence or presence of various concentrations of cisplatin (**Figure 4B**). We found that keratin fusion variant-expressing cells exhibited a higher survival advantage than control cells (**Figure 4B**). The higher cisplatin concentration, the higher survival proportion was found in keratin fusion variant expressing cells, especially in giant-like KF expressing ones (**Figure 4B**). It implies keratin fusion variant plays a critical role in drug resistance for tumor recurrence.

**Figure 4.**
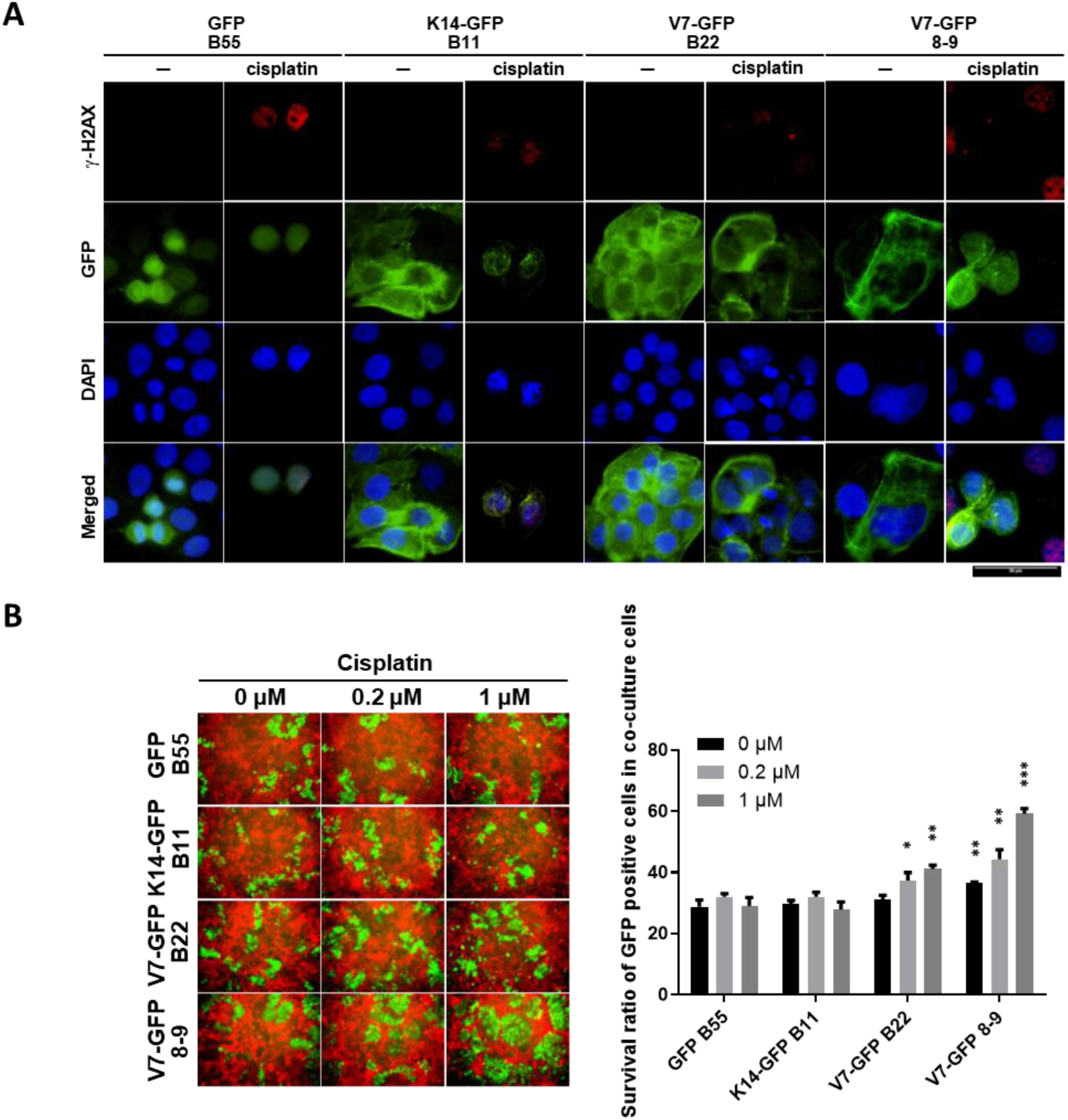
Keratin fusion variant withstands chemotherapy agent treatment. (A) Keratin expression sublines cells were treated with 10 μM cisplatin for 24 hr or not, and followed by performing immunofluorescence against γ-H2AX and nuclei visualization with DAPI staining. (B) Keratin expression sublines cells (5×10^3^) were co-cultured with CAL27 cells harboring mCherry expression (2×10^4^) as a control group on a 12-well plate and treated with the indicated concentration of cisplatin for 6 days. To observe the respective cells with a fluorescence microscope (OLYMPUS IX83). Percentages of cell viability in keratin-expressing CAL27 cells post-treating with cisplatin for 6 days were determined by calculating the occupied areas of indicated cells. The occupied areas were measured by Image J software. *P < 0.05, **P < 0.01 and ***P < 0.001 according to a Student’s t-test.

### Keratin fusion variant promotes cell motility

Keratin fusions in OSCC are correlated with poor recurrence/ lymph node metastasis-free survival [7]. Furthermore, we found enhanced activity for the regulation of actin cytoskeleton and the regulation of actin filament bundle assembly in KFs expressing cells (**Figure 5A**). Therefore, we determine whether the KF variant influences actin cytoskeleton organization by staining filament actin with phalloidin (**Figure 5B**). We found that KF variant-expressing cells displayed higher intensity and length of F-actin (**Figure 5C**). Actin network organization is involved in cell motility. Therefore, we determined whether the KF variant affected cell motility by using an *in vitro* migration and invasion assay. We found that KF variant expressing cells harbored higher cell motility activity (**Figure 5D and 5E**).

**Figure 5.**
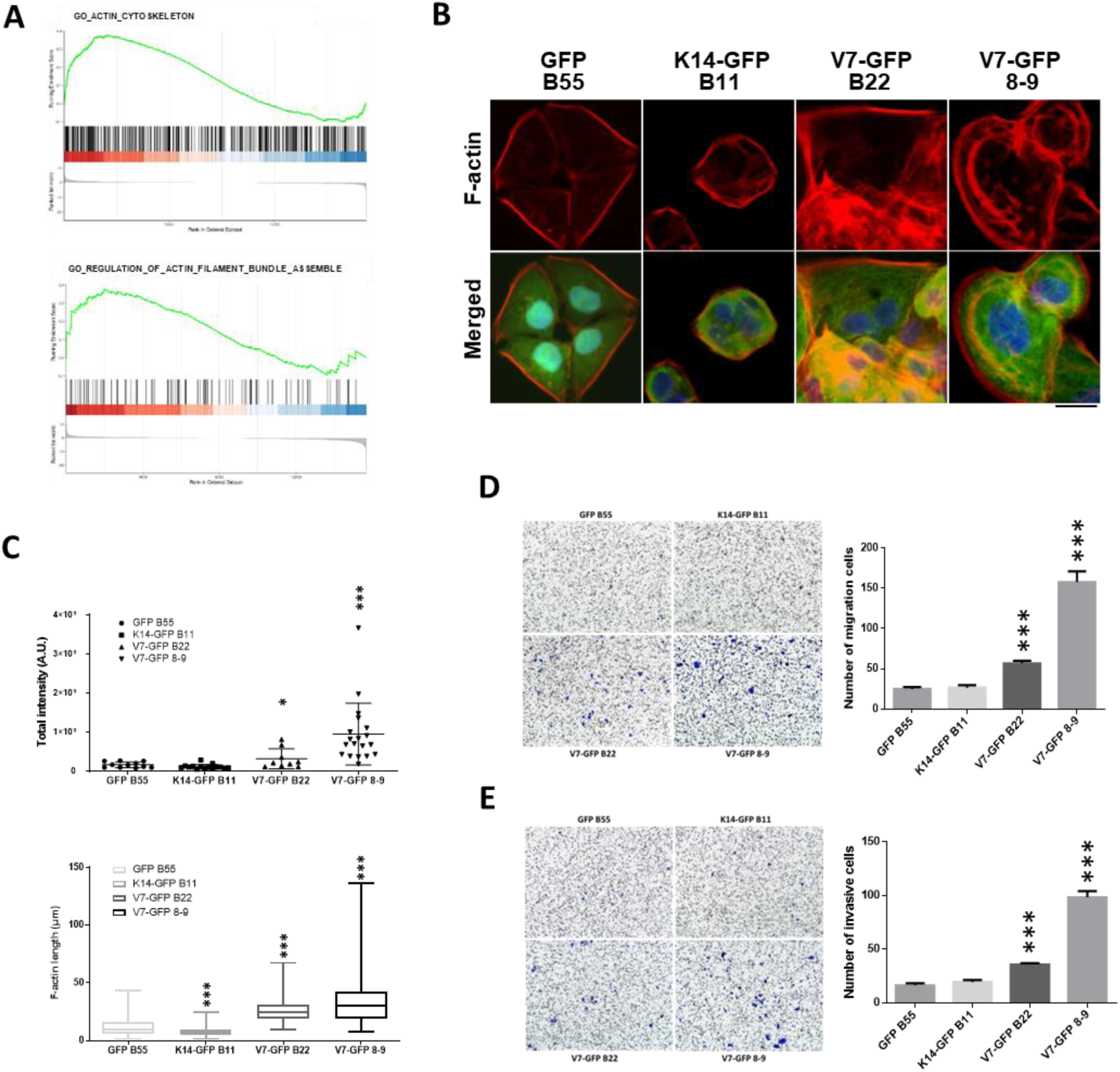
Keratin fusion mutation is involved in cytoskeleton remodeling and enhances cell motility. (A) Gene Ontology enrichment analysis was performed to reveal the enhanced activity in the regulation of actin cytoskeleton between the control cells and keratin fusion expressing ones. (B) Keratin-expressing cell sublines were fixed and stained with Rhodamine Phalloidin to visualize the network of filamentous actin. Nuclei in cells were visualized with DAPI staining. Scale bar indicates 20 μm. (C) The total intensities (upper panel) and lengths of actin filaments (bottom panel) per cell were analyzed with CellSens software. (D) The cell migration activity of keratin-expressing cells was measured by using the transwell assay followed by crystal violet staining. Briefly, 5 × 10^4^ cells suspended in serum-free medium were seeded onto the upper chamber of the transwell, and the complete medium with 10% FBS was loaded onto the lower chamber. After incubation for 24 hours at 37°C, the migrating cells were harvested and stained with crystal violet solution for analysis. The numbers of migration cells by crystal violet staining were counted under a fluorescence microscope at magnification × 100. We randomly selected five fields to count the number of invasive cells. The data represent the means ±SD from 3 separate experiments. ***P < 0.001 based on Student’s t-test. (E) The cell invasion activity of keratin-expressing cells was measured by using Matrigel-coated transwell inserts followed by crystal violet staining. The procedure mentioned above in (D), except that the transwell inserts were coated with 10% Matrigel. The data represent the means ±SD from 3 separate experiments. ***P < 0.001 based on Student’s t-test.

### Enforcement of cell cycle progression-associated signaling drives PACC formation in keratin fusion variant-expressing cells

From experiments mentioned above, the keratin fusion variant promotes genomic instability but withstands various stresses for survival via distinctive mechanisms to control cell fate. Giant cell-like KF cell sublines harbored higher expression of cell cycle-associated and DNA damage response-associated molecules and suppressed cGAS-STING-mediated cell death signaling. Here, we wanted to decipher which signaling axes contribute to PACC formation and aggressiveness. 607 DEGs (with more than fourfold increase) were found in 8-9 subline. These DEGs were subjected to STRING analysis [44]. We found that three major cluster genes were enriched in 8-9 sublines. The three ones are composed of 1) growth factors and receptors; 2) mitochondria-associated molecules, and 3) cell cycle-associated ones (**Figure 6A**). We further wanted to identify the corresponding upstream transcription factors for the 607 DEGs to drive PACC formation by ChEA3 analysis [45]. The top 30 transcription factors and involved gene counts were plotted in a scatter diagram, and the top 3 critical factors were labelled (**Figure 6B**). The top 10 transcription factors (**Table 1**) are involved in cell cycle progression.

**Figure 6.**
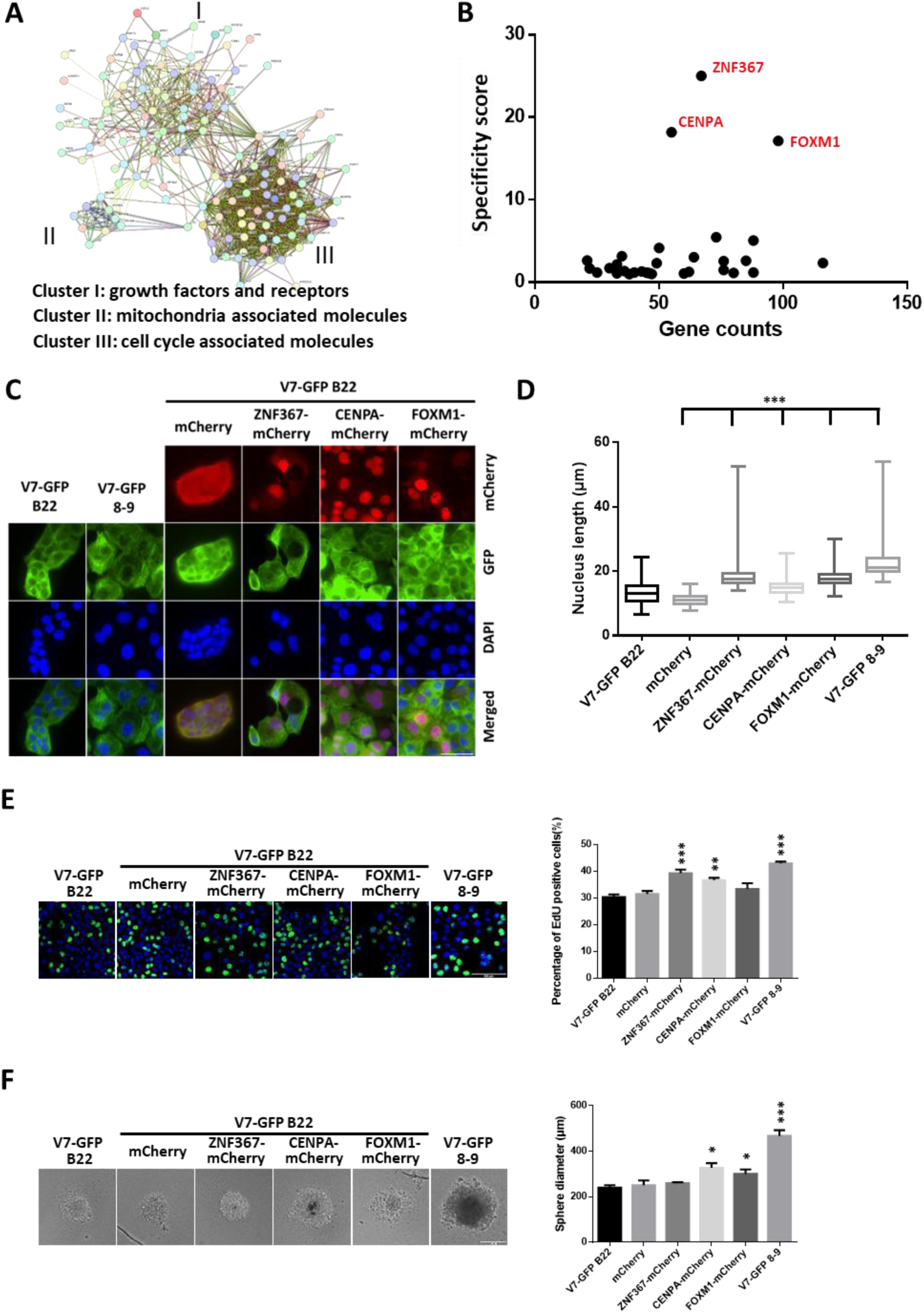
Enforcement of ZNF367, CENPA, or FOXM1 signaling drives PACC formation in keratin fusion variant-expressing cells. (A) The upregulated genes (n=607) in V7-GFP 8-9 cells (with more than 4-fold of expression than B22 cells) were subjected to STRING analysis, and the three major cluster genes were enriched and plotted. (B) The DEGs were subjected to ChEA3 analysis to identify the corresponding transcription factors to drive PACC formation (Table 1). The top 30 transcription factors and involved gene counts were plotted in a scatter diagram, and the top 3 critical factors were labelled. (C) mCherry-fusion proteins expressing cell sublines were fixed and stained with against mCherry antibody to visualize the expression of mCherry-fusion proteins. Nuclei in cells were visualized with DAPI staining. Scale bar indicates 50 μm. (D) The lengths of nuclei were measured and plotted in a box plot. (E) EdU labelling assay was used to determine the ratio of S phase in keratin fusion variant expressing sublines. Scale bar indicates 100 μm. (F) Sphere formation assay was used to determine the cancer stem characteristics in keratin fusion variant expressing sublines. Scale bar indicates 200 μm. (G) The keratin-expressing cell lines were harvested and lysed. The lysates were subjected to immunoblotting for the indicated proteins. The expression levels of β-actin were used as loading controls. The data represent the means ±SD from 3 separate experiments. *P < 0.05, **P < 0.01, ***P < 0.001 based on Student’s t-test.

**Table 1.** Enforced upstream signaling networks in the KF giant cell subline^a^. ^a^A total of 607 differentially expressed genes (genes with more increase 4 folds in 8-9 sublines as compared to B22 cells) were subjected to CHEA3 analysis [45] to identify the critical upstream transcription factors for the induction of the corresponding differentially expressed genes leading to PACC formation.

| Rank | TF | Score | Overlapping Genes |
| --- | --- | --- | --- |
| 1 | **ZNF367** | 4 | 67 |
| 2 | **CENPA** | 5.5 | 55 |
| 3 | **FOXM1** | 5.833 | 98 |
| 4 | **HMGA2** | 18.33 | 73 |
| 5 | **E2F7** | 19.8 | 88 |
| 6 | **DNMT1** | 24 | 50 |
| 7 | **ZNF878** | 31.5 | 35 |
| 8 | **ZNF93** | 33 | 64 |
| 9 | **ZNF888** | 38 | 21 |
| 10 | **TFDP1** | 38.25 | 85 |

We further determined whether upregulation of ZNF367, CENPA, or FOXM1 contributes to PACC formation in V7-GFP B22 cells. We used the IF strategy to determine the expression of the indicated transcription factors due to the low expression of mCherry-fusion proteins in V7-GFP B22 cells. We found that upregulation of ZNF367, CENPA, or FOXM1 promotes polyaneuplody formation (**Figure 6C**) and increases nuclei lengths compared to the control cells (mCherry expressing V7-GFP B22 ones) (**Figure 6D**). From previous results (**Figure 1C**), giant cell-like KF expressing cells (8-9 subline) harbored a higher ratio of S phase than B22 cells. We also determined whether upregulation of ZNF367, CENPA, or FOXM1 contributes to cell cycle progression in V7-GFP B22 cells. We adopted the EdU labelling assay to measure the active replication phase in the expressing indicated transcription factor cell sublines. We found that upregulation of ZNF367, CENPA, or FOXM1 dramatically increased the ratio of EdU-positive cells (**Figure 6E**). In addition to cell proliferation, we also determined whether upregulation of ZNF367, CENPA, or FOXM1 dramatically advances the cancer stemness to drive aggressiveness by using the sphere formation assay. We found that giant cell-like V7 variant expressing cells harbored dramatically higher stemness activity than B22 cells (p=0.0001) (**Figure 6F**). Especially, CENPA or FOXM1 overexpression can significantly increase the sphere diameter (p=0.0112; p=0.0369). In addition to the impacts of transcription factors on cell abilities of cell proliferation and stemness, we also evaluated the impact of the corresponding transcription factors on clinical significance in head and neck cancer patients. The results showed that ZNF367, CENPA, and FOXM1 play critical roles in aneuploidy and hypoxia with grades in head and neck cancer patients (**Figure S4**). It implies that these transcription factors contribute to tumor aggressiveness via multiple perspectives.

## Discussion

Keratin intermediate filament is a critical component of the cytoskeleton network, which establishes the necessary mechanical support to withstand environmental stresses. Dysregulation of intermediate filaments causes diseases associated with dermatology and even cancers. KFs are correlated with poor prognosis via enhancing cancer cell stemness and cancer aggressiveness [7]. In the present study, we revealed that keratin fusion variant expression promotes genomic instability through loss of nuclear membrane integrity, alteration of DNA damage responses, and cytokinesis defects-mediated centrosome amplification. Maheswaran’s team found that TGF-β-induced EMT results in cytokinesis failure, leading to genomic instability [46]. Our previous study also revealed that KFs expression upregulates TGF-β signaling via protein array analyses, which might be a partial contribution to genomic instability [7]. Genomic instability is a critical cancer hallmark [10] leading to the acquisition of biological capabilities required for tumorigenesis and progression [47]. An increasing body of evidence suggests that genomic instability drives drug resistance and metastasis [30, 48–50]. Besides, genomic instability is also the critical driving force for intratumor heterogeneity and tumor evolution [51, 52]. In the meantime, we showed that keratin fusion variant promoted chemotherapy agent resistance and enhanced cell motility, which contributes to tumor aggressiveness in the present study.

In the present study, we showed that KF variant expression promotes genomic instability accompanied by micronucleus formation (**Figure 1**). cGAS-STING-mediated type I interferon responses were responsible for targeting cells containing cytoplasmic micronuclei (MN) to eliminate chromosome instability (CIN) [24, 32]. Bruyn’s group revealed that IL-6/IL-6R regulates STAT3 signaling to enhance cell survival and growth of cancer cells harboring cDNA-mediated cGAS-STING signaling [35]. We found one of the KF variant clones activates cGAS-STING-mediated STAT1 activation for type 1 interferon response, but also activates STAT3 signal for cell survival and growth upon cGAS-STING signaling. In the meantime, giant cell-like cells harboring KF variant cells suppress cGAS-STING signaling and expression (**Figure 3D**). Positive feedback regulation between cGAS-STING signaling and type I IFN was demonstrated [53, 54]. Interestingly, giant cell-like cells harboring KF variant cells showed reduced levels of cGAS, STING, and STAT1 (**Figure 3D**). It indicates dramatically suppressing cGAS-STING mediated cell death signaling. From Liu’s study [55], ECM stiffness-involved mechanotransduction can modulate cGAS-STING signaling. We found that keratin fusion variant expressing cells harbored higher levels of F-actin network contributing to cell motility and mechanotransduction (**Figure 5**). Therefore, Keratin fusion variant expressing cells affect cGAS-STING signaling for cancer cell survival.

To further characterize which signaling cascades contribute to this cell fate, we analyzed the differentially expressed genes and their corresponding upstream transcription factors by ChEA3. We individually enforced expression of the transcription factors in V7-GFP B22 cells to determine whether overexpression of the specific factor can switch B22 cells to giant-like cells. We found that upregulation of ZNF367, CENPA, and FOXM1 obviously increased the nucleus size and proliferation ratio in V7-GFP B22 cells (**Figure 6C and D**). It indicates that upregulation of these factors contributes to genomic instability and increased cell cycle progression. However, upregulation of these factors in V7-GFP B22 cells could not suppress cGAS-STING signaling (**Figure 6E**). It might imply that the expression levels of transcription factors are not enough to pass the threshold, or a single factor is not sufficient to trigger the switch to aggressive and giant ones. But ZNF367, CENPA, and FOXM1 are still feasible therapeutic targets for cancer treatment [56–60].

Dysregulation of keratin not only influences the intermediate filament network but also affects the whole cytoskeleton network. In our previous study, the keratin fusion variant would perturb the distribution of other endogenous keratin isoforms [7]. In the present study, we found that keratin fusion variant expression promotes the reorganization of the actin network via strengthening F-actin distribution, leading to increased cancer cell motility (**Figure 5**). In addition to the actin network, the keratin fusion variant also influences the microtubule network. Centrosome is known as the primary microtubule-organizing center to control the microtubule network [38]. In the present work, we found that keratin fusion variant promotes genomic instability partly through cytokinesis defects caused by centrosome amplification (**Figure 1**). We also showed that the keratin fusion variant altered the expression levels of centriole biogenesis and cytokinesis associated genes (**Figure 3**). Briefly, keratin dysregulation may affect the whole cytoskeleton network and contribute to the alteration of cell fates and tumor progression [2].

In conclusion, our data reveal that the keratin fusion acts as an independent prognostic factor for tumor aggressiveness through not only increased cancer stemness but also enhanced genomic instability. Genomic instability promotes PACC formation, drug resistance, suppression of cGAS-STING signaling, and tumor evolution, leading to tumor aggressiveness (**Figure 7**).

**Figure 7.**
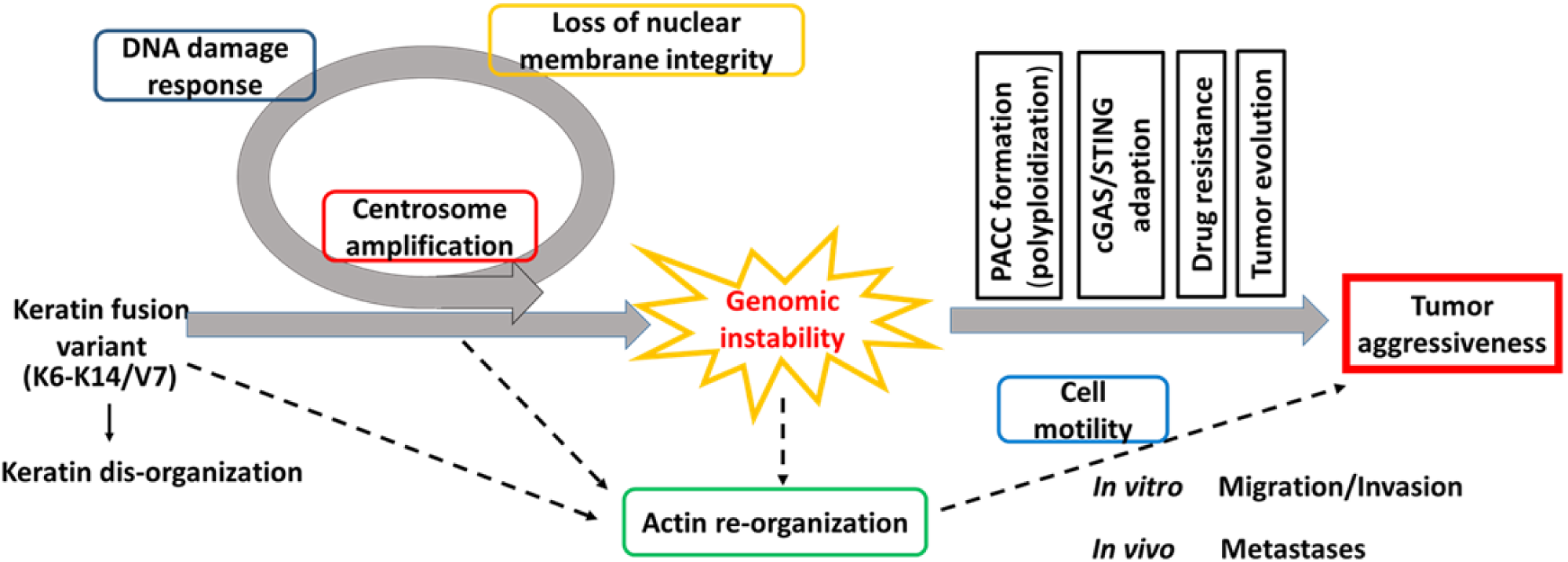
Working hypothesis for keratin dysregulation-driving tumor aggressiveness. Keratin fusion promotes genomic instability through loss of nuclear membrane integrity, alteration of DNA damage response signaling, and induction of centrosome amplification, accompanied by actin network reorganization. Genomic instability triggers polyaneuploid cancer cell formation, cGAS-STING adaptation, drug resistance, and tumor evolution to exacerbate tumor malignancy.

## Author contribution

HHC, CYYC, and JJCS provided the study concept and design. HHC, BYTK, KS, LYTT, TH, and CMC performed the experiments. HHC, BYTK, KS, YCC, PKB, and MSL interpreted and analyzed the data. HHC and PKB wrote the original draft. HHC, CYYC, and JJCS revised and edited the manuscript. All authors have read and agreed to the final version of the manuscript.

## Institutional Review Board Statement

The present study was conducted in accordance with the Declaration of Helsinki, and approved by the Institutional Review Board of China Medical University Hospital (**CMUH102-REC1-009**).

## Acknowledgments

We thank Inflammation Core Facility at Academia Sinica, Taiwan for technical support. The core facility is funded by the Academia Sinica Core Facility and Innovative Instrument Project (AS-CFII-113-A9).

